# Small-scale spatial structure influences large-scale invasion rates

**DOI:** 10.1101/814582

**Authors:** Michael J. Plank, Matthew J. Simpson, Rachelle N. Binny

**Affiliations:** School of Mathematics and Statistics, University of Canterbury, Christchurch, New Zealand; Te Pūnaha Matatini, Centre of Research Excellence, New Zealand; School of Mathematical Sciences, Queensland University of Technology, Brisbane, Australia; Manaaki Whenua, Lincoln, New Zealand

**Keywords:** density-dependence, dispersal, mean-field model, plant populations, species range shifts, stochastic model

## Abstract

Local interactions among individual members of a population can generate intricate small-scale spatial structure, which can strongly influence population dynamics. The two-way interplay between local interactions and population dynamics is well understood in the relatively simple case where the population occupies a fixed domain with a uniform average density. However, the situation where the average population density is spatially varying is less well understood. This situation includes ecologically important scenarios such as species invasions, range shifts, and moving population fronts. Here, we investigate the dynamics of the spatial stochastic logistic model in a scenario where an initially confined population subsequently invades new, previously unoccupied territory. This simple model combines density-independent proliferation with dispersal, and density-dependent mortality via competition with other members of the population. We show that, depending on the spatial scales of dispersal and competition, either a clustered or a regular spatial structure develops over time within the invading population. In the short-range dispersal case, the invasion speed is significantly lower than standard predictions of the mean-field model. We conclude that mean-field models, even when they account for non-local processes such as dispersal and competition, can give misleading predictions for the speed of a moving invasion front.

## Introduction

Spatial structure can affect population dynamics. Common examples of spatial structure are clustering, where individuals tend to occur in tightly packed groups, and regular structure, where individuals tend to be evenly spaced from one another (Pacala and Silander Jr, 1985; Mahdi and Law, 1987; Purves and Law, 2002). Spatial structure can arise from individual-level processes and interactions that occur locally in space, such as competition (Yokozawa et al., 1999; Adams et al., 2013), dispersal (Lewis and Pacala, 2000), adhesion (Johnston et al., 2013) and crowding (Binny et al., 2016a). These local interactions typically generate small-scale spatial structure that occurs at a length scale of the order one to ten times an individual’s size. Despite being local in origin, spatial structure can have significant large-scale effects on population size, and even determine whether the population survives or dies (Law et al., 2003). Mean-field models neglect short-range correlations among individual locations by assuming that the presence of an individual at one location is independent of the presence of an individual at neighbouring locations. A consequence of this assumption is that individuals interact with one another in proportion to their average densities. Mean-field models include spatially explicit models, such as partial differential equations and integro-differential equations (Hastings et al., 2005), which describe large-scale spatial variations in average density, but cannot account for the effects of small-scale spatial structure.

Most mathematical studies that incorporate small-scale spatial structure in a lattice-free setting have focused on the relatively simple case where the population occupies a fixed region at constant average density (e.g. Bolker and Pacala, 1999; Dieckmann et al., 2000; Murrell and Law, 2003; Murrell, 2005; Binny et al., 2016b). This does not preclude the development of spatial structure over time. For example, individuals may become clustered and so, in any given realisation of the process, the density will be higher in some regions than others. However, the locations of the clusters are random and, when averaged over multiple realisations, the density is spatially uniform. We refer to this as the *translationally invariant* case.

The spatial stochastic logistic model (Bolker and Pacala, 1997; Law et al., 2003) is a spatially explicit, individual-based model of dispersal and competition that is translationally invariant. In this model, individuals undergo density-independent proliferation accompanied by dispersal, and density-dependent mortality, with the mortality rate being an increasing function of the number and the proximity of individuals within the local neighbourhood. The mean-field equation for the spatial stochastic logistic model is the well-known logistic growth differential equation (Law et al., 2003). However, depending on the spatial scales of dispersal and competition, the stochastic model can produce different population dynamics to the mean-field equation, in both its transient and its long-term phase. An improvement on the mean-field model can be obtained using spatial moment dynamics (Dieckmann and Law, 2000; Plank and Law, 2015) to account for the pair density function (second spatial moment) as well as the average density (first spatial moment). Law et al. (2003) used this approach to show that, when there is a regular spatial structure, the population grows to a higher density than predicted by the mean-field equation. When there is a clustered structure, the population eventually asymptotes to a lower density than predicted by the mean-field equation, or can even die out altogether (Law et al., 2003). Translationally invariant models such as these can investigate the effect of spatial structure on population density, but cannot describe populations where the occupied region changes over time. These models are therefore not suitable for modelling many ecological scenarios such as species invasions, range shifts, or moving population fronts.

Some studies have investigated the more complex *translationally dependent* case, where the region occupied by the population changes over time. Lewis and Pacala (2000) focused on a model of invasion with density-independent proliferation and long-range dispersal, and used a spatial moments approach to derive results for the dependence of invasion speed on the dispersal kernel. Lewis (2000) generalised this model to show that local density-dependent proliferation reduced the invasion speed in one spatial dimension, relative to mean-field predictions. However, this model only applied to a situation where density-dependence affects proliferation, and operates over a short spatial scale. Omelyan and Kozitsky (2019) derived a spatial moments approximation for the translationally dependent version of the spatial stochastic logistic model in one dimension. They showed that the results differed significantly from the mean-field model, which neglects correlations among individual locations. However, they did not test the predictions of their spatial moment equations against individual-based simulations, which serve as ‘ground truth’ for the approximation. It is therefore unknown how well the spatial moment dynamics system approximates the underlying stochastic process in practice.

Invasion dynamics with small-scale spatial structure have also been studied in lattice-based models. Ellner et al. (1998) developed a method for estimating invasion speed using a pair-edge approximation, which is a type of pair density function for a lattice-based model. Simpson and Baker (2011) used a pair approximation to improve mean-field predictions for the dynamics of moving fronts. A key drawback of lattice-based models is that they limit the types of interactions and spatial structure that the model can support (Plank and Simpson, 2012). In the models referred to above, dispersal or movement is restricted to nearest-neighbour lattice sites and competition or crowding is effectively modelled by volume exclusion, i.e. a maximum of one individual is allowed per lattice site. This restricts the scope for investigating the interplay between these individual-level mechanisms as their spatial scales vary. In addition, the geometry of the lattice can impose an artificial carrying capacity (Plank and Simpson, 2012), affect invasion dynamics (Fernando et al., 2010), and may have other unknown consequences.

In this study, we investigate the dynamics of an invading population with small-scale spatial structure, using the translationally dependent version of the spatial stochastic logistic model. Similarly to Omelyan and Kozitsky (2019), we focus on the case where the population is initially confined to a subregion of the domain and subsequently invades via dispersal of individuals into previously unoccupied regions. However, unlike Omelyan and Kozitsky (2019), we carry out individual-based simulations of the translationally dependent spatial stochastic logistic model and we work in a two-dimensional domain. We systematically investigate scenarios with different spatial scales for competition and dispersal, and compare them to predictions of the mean-field model. This allows us to quantify the departure of the stochastic process from mean-field dynamics in terms of the spatial structure. To provide insight into how spatial structure affects population spreading, we test how the speed of the invasion and the population density behind the invasion front depend on the spatial scales of competition and dispersal. We interpret these results in light of what is already known about the translationally invariant form of the spatial stochastic logistic model.

## Translationally dependent spatial stochastic logistic model

### Individual-based model

We consider a population of *N*(*t*) individuals with locations *z*_*i*_(*t*) = (*x*_*i*_(*t*), *y*_*i*_(*t*)) ∈ Ω ⊆ ℝ^2^ (*i* = 1, …, *N*(*t*)). The spatial stochastic logistic model consists of two individual-level mechanisms: density-independent proliferation accompanied by dispersal; and density-dependent mortality modelling local competition. Specifically, in a short time interval *δt*, each of the *N*(*t*) agents has a probability λ*δt* + *O*(*δt*^2^) of proliferating, independent of all other agents. Offspring are dispersed to a location at a displacement *ξ* from the parent, where *ξ* is a random variable from a bivariate probability distribution with density function *w*_*d*_(*ξ*), referred to as the dispersal kernel. In addition, agent *i* has a probability *μ*_*i*_(*t*)*δt* + *O*(*δt*^2^) of dying in time interval *δt*. The mortality rate for individual *i* at time *t* consists of a constant density-independent term *μ*_0_ and a contribution from neighbouring individuals *μ_c_*, weighted by a competition kernel *w*_*c*_(*ξ*):

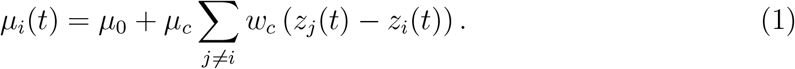

The dispersal and competition kernels are assumed to be isotropic and symmetric about the origin and to integrate to 1 over Ω. We consider a rectangular domain Ω = [−*L*_*x*_, *L*_*x*_]× [0, *L*_*y*_] with periodic boundaries, such that dispersal and competition are wrapped across opposing boundaries. This is equivalent to the spatial stochastic logistic model studied by Bolker and Pacala (1997) and Law et al. (2003) for a translationally invariant population. To investigate the dynamics of a translationally dependent population, we consider an initial condition where *N*_0_ agents are distributed independently and uniformly at random in the region −*x*_0_ ≤ *x* ≤ *x*_0_, where *x*_0_ < *L*_*x*_.

Since the population is translationally invariant in the vertical direction, we calculate the average agent density 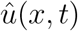 in thin vertical strips of width *δx*:

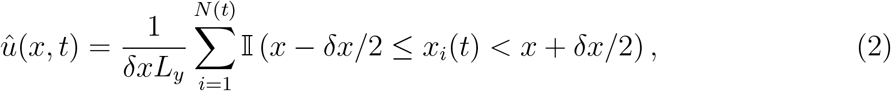

where 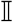 is an indicator function.

To quantify the spatial structure of the population, we compute the pair correlation function at time *t* defined by:

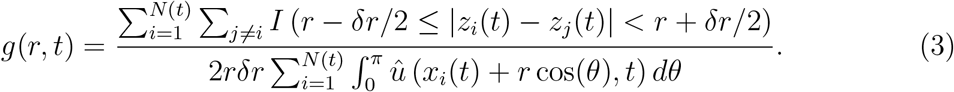

This corresponds to the ratio of the number of pairs a distance *r* apart to the expected number of pairs a distance *r* apart, in a population with density 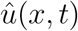 that is in a state of complete spatial randomness. The integral in Eq. (3) is approximated numerically by discretising the integration variable *θ* and using linear interpolation for the required values of 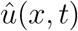. In principle, the nature and strength of spatial structure could vary across the spatial domain Ω. This could be measured by calculating different pair correlation functions in different regions *R* ⊂ Ω by restricting the index *i* to individuals that are in the region *R*. However, in practice we find that the pair correlation function is very similar throughout Ω, so for simplicity we calculate a single pair correlation function across the whole of the spatial domain.

We measure the extent of the invasion at time *t* by calculating the mean squared displacement, defined as the average value of *x*_*i*_(*t*)^2^ across all *N*(*t*) agents. We also measure the location of the invasion front at time *t* as the location of the agent with the 10^th^ largest value of |*x*_*i*_(*t*)|. We use the 10^th^ largest value as opposed to the largest value to reduce noise caused by outlying agents, but the qualitative results are not sensitive to this choice.

We perform *M* independently initialised realisations of the individual-based model (IBM) and average 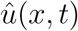 and *g*(*r, t*) over the *M* realisations. Note that these are effectively double averages because Eq. (2) is the average density in a thin vertical strip and Eq. (3) is the average pair density in a thin annulus. This means that smooth outputs can typically be obtained with a relatively small number of IBM realisations. The dispersal and competition kernels are set to be bivariate Heaviside functions:

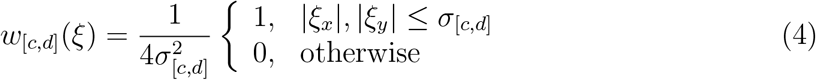

This provides a simple on/off model for spatial interactions in which competition and dispersal are uniform in a specified neighbourhood (a square of length *σ*_*c*_ and *σ*_*d*_ respectively) and zero outside that neighbourhood. We also test the effect of using Gaussian functions instead of Heaviside functions for the interaction kernels (see Supplementary Information). Parameter values are shown in Table 1.

**Table 1:** Model parameter values.

| Parameter | Value |
| --- | --- |
| Proliferation rate | $\lambda = 1$ |
| Intrinsic mortality rate | $\mu_0 = 0.01$ |
| Neighbour-dependent mortality rate | $\mu_c = 0.1$ |
| Dispersal kernel standard deviation | $\sigma_d$ (variable) |
| Competition kernel standard deviation | $\sigma_c$ (variable) |
| Domain size | $L_x = 20, L_y = 10$ |
| Width of the initially occupied region | $x_0 = 1$ |
| Initial population size | $N_0 = 20$ |
| Threshold density for invasion front | $u_{\text{thresh}} = 1$ |

### Mean-field dynamics

The mean-field equation for the translationally dependent spatial stochastic logistic model is:

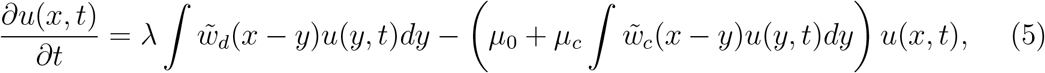

where *x* ∈ [−*L*_*x*_, *L*_*x*_]. This formulation makes use of the translational invariance in the vertical direction to write the average population density *u*(*x, t*) in terms of the horizontal coordinate *x* only, where 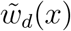 and 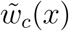 are the marginal distributions over *x* of the dispersal and competition kernels *w*_*d*_(*x, y*) and *w*_*c*_(*x, y*) respectively. Eq. (5) neglects correlations in the locations of pairs of agents and assumes that the system is locally well mixed. Formally, this corresponds to approximating the joint density of pairs of agents at *x* and *y* in the second integral in Eq. (5) by the product of the average agent densities *u*(*x, t*) and *u*(*y, t*).

The population carrying capacity *K* (i.e. equilibrium average density in a uniformly occupied domain) can be found from Eq. (5). This corresponds to a solution *u* to Eq. (5) that is independent of both *x* and *t*, which is uniquely given by *K* = (λ − *μ*_0_)/*μ*_*c*_. We can also calculate the total population size *N*(*t*) and mean squared displacement *MSD*(*t*) at time *t* under the mean-field equation via:

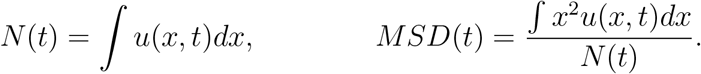

The location of the invasion front at time *t* is defined to be the smallest value of |*x*| for which *u*(*x, t*) > *u*_thresh_.

The integro-differential equation (5) is solved by discretising *x* using a mesh spacing *δx* = 0.01 and solving the resulting system of ordinary differential equations using Matlab’s *ode45* routine. To implement periodic boundaries, we set 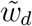 and 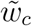 to be periodic extensions of the dispersal kernel and competition kernel respectively on *x* ∈ [−*L*_*x*_, *L*_*x*_]. This means that population members located near the boundary at *x* = −*L*_*x*_ are interacting with population members located near the boundary at *x* = *L*_*x*_ and vice versa.

## Results

First, we test the behaviour of the translationally dependent spatial stochastic logistic model when both dispersal and competition operate over a long range (*σ*_*d*_ = *σ*_*c*_ = 5, Fig. 1). In this case, agents compete weakly with neighbours over a relatively large neighbourhood (encompassing the full height *L*_*y*_ of the domain), and the correlation between locations of parent and offspring is weak. As a consequence, spatial structure is close to random (pair correlation function close to 1, Fig. 1c) and the IBM results are close to the predictions of the mean-field equation (Fig. 1d-e).

**Figure 1:**
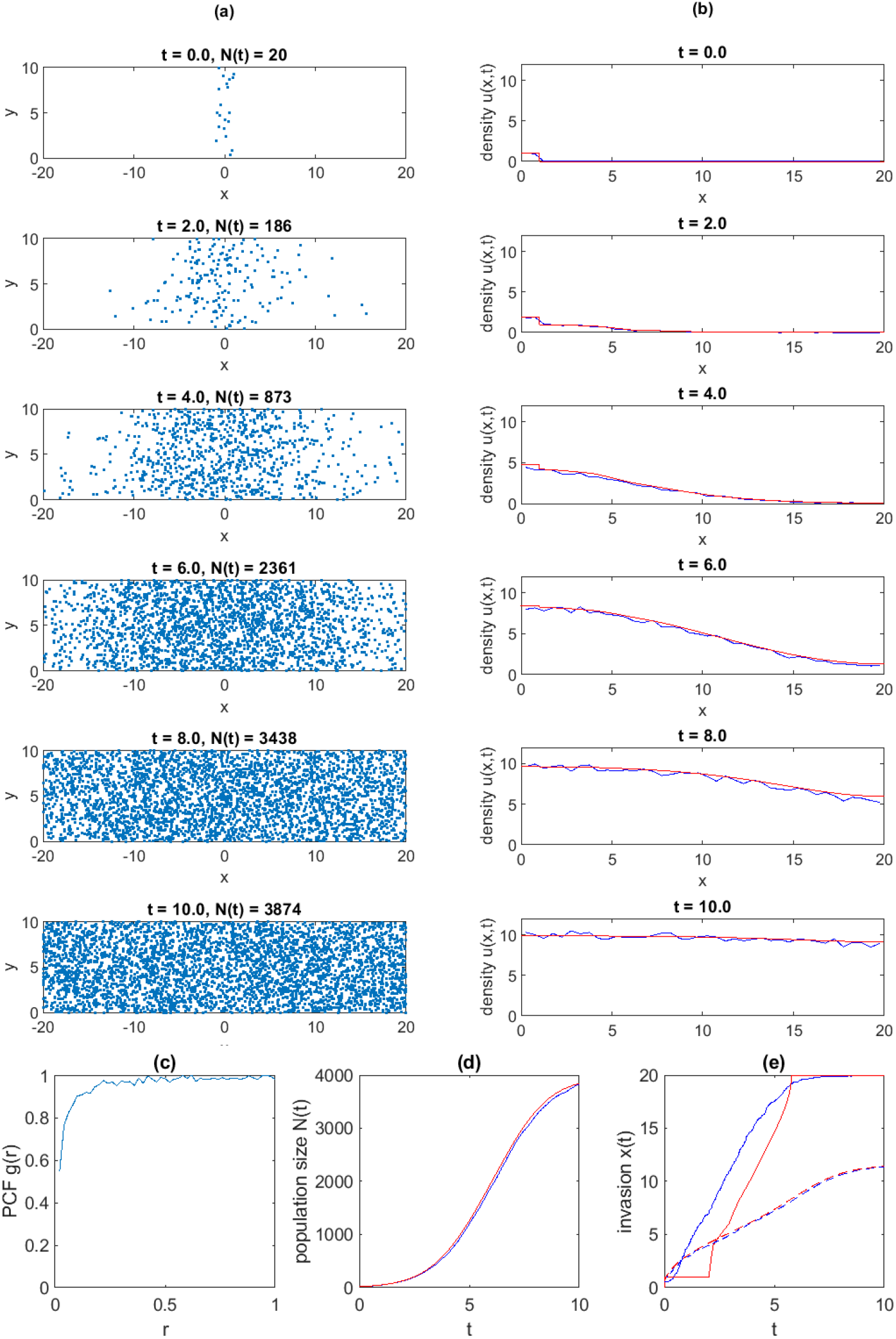
IBM and mean-field results for long-range competition and long-range dispersal (*σ*_*c*_ = 5, *σ*_*d*_ = 5): (a) snapshots of a single realisation of the IBM; (b) average agent density in the IBM (blue) and mean-field equation (red) at *t* = 0, 2, 4, 6, 8, 10. (c) pair-correlation function (PCF) at *t* = 10; (d) time series of the average population size in the IBM (blue) and mean-field equation (red); (e) time series of the invasion size measured by the location of the invasion front (solid) and the root mean squared displacement (dashed) in the IBM (blue) and mean-field equation (red). IBM results in (b-e) are averaged across *M* = 10 independent realisations, each initialised with *N*_0_ agents randomly placed in the region |*x*| < *x*_0_.

We now focus on the deviation from mean-field dynamics as the range for dispersal *σ*_*d*_ and/or competition *σ*_*c*_ are reduced. In all cases, the long-term statistical equilibrium of the model is a spatially structured population with uniform average agent density, consistent with the translationally invariant version of the spatial stochastic logistic model (Law et al., 2003). Here, we focus on the population dynamics in the transient phase corresponding to the invasion of the initially unoccupied region.

When competition is short-range and dispersal is long-range (*σ*_*c*_ = 0.1, *σ*_*d*_ = 1, Figure 2), a regular spatial structure develops, indicated by values of the pair correlation function less than 1 for pairs less than distance 0.1 apart (Fig. 2c). These results are consistent with the translationally invariant spatial stochastic logistic model: strong competition in small neighbourhoods makes the probability of more than one agent persisting in such a neighbourhood very small. Conversely, offspring have a high probability of escaping the competitive influence of their parent and finding an empty neighbourhood. The population in the region behind the invasion front reaches a substantially higher density then predicted by the mean-field equation (Fig. 2b,d) because a typical agent experiences a lower-density neighbourhood, and therefore a lower mortality rate, than in the mean-field model. However, the speed of the invasion, as measured by the root mean squared displacement or by the location of the invasion front, is well predicted by the mean-field equation (Fig. 2e).

**Figure 2:**
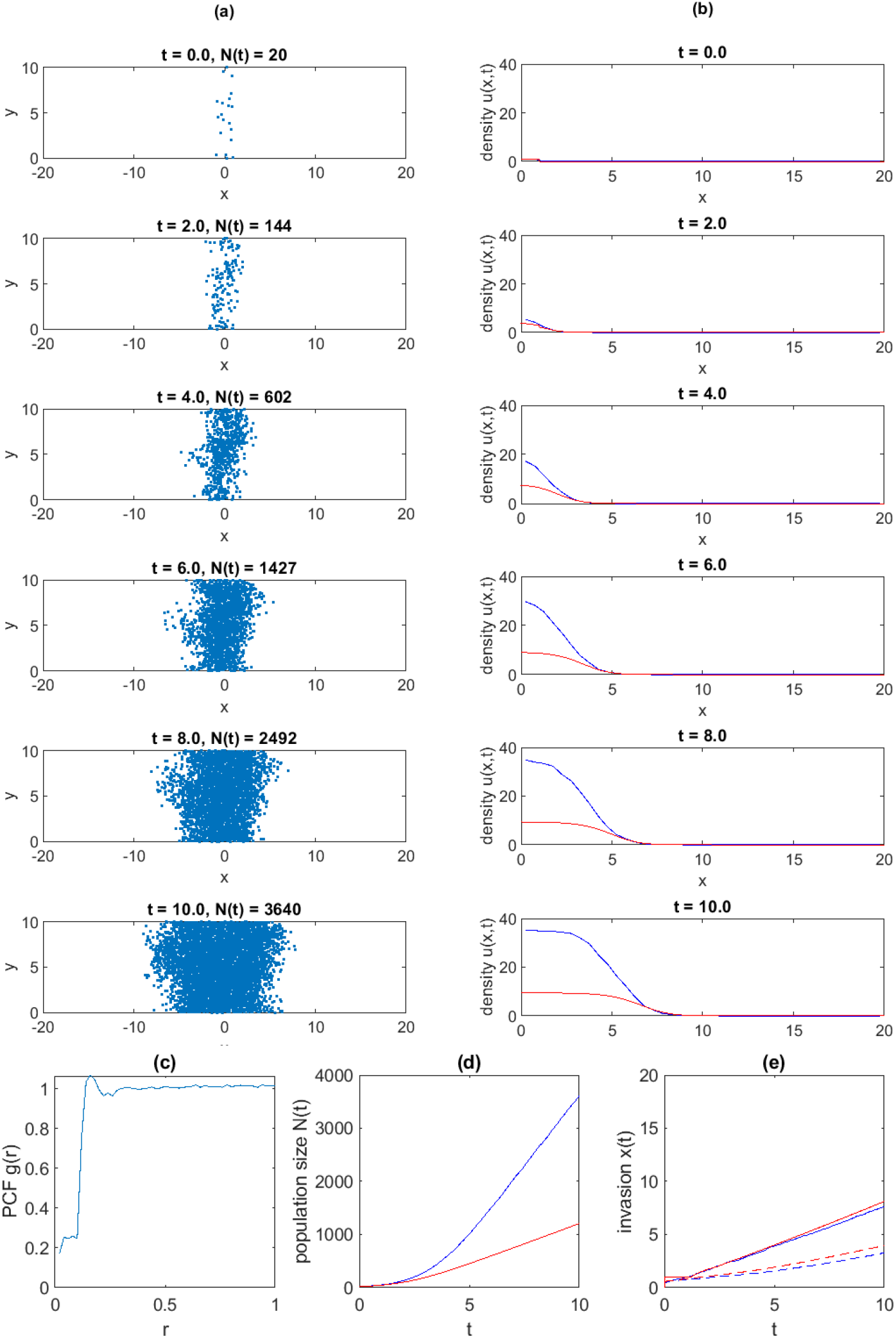
IBM and mean-field results for short-range competition and long-range dispersal (*σ*_*c*_ = 0.1, *σ*_*d*_ = 1): (a) snapshots of a single realisation of the IBM; (b) average agent density in the IBM (blue) and mean-field equation (red) at *t* = 0, 2, 4, 6, 8, 10. (c) pair-correlation function (PCF) at *t* = 10; (d) time series of the average population size in the IBM (blue) and mean-field equation (red); (e) time series of the invasion size measured by the location of the invasion front (solid) and the root mean squared displacement (dashed) in the IBM (blue) and mean-field equation (red). IBM results in (b-e) are averaged across *M* = 10 independent realisations, each initialised with *N*_0_ agents randomly placed in the region |*x*| < *x*_0_.

When competition is long-range and dispersal is short-range (*σ*_*c*_ = 1, *σ*_*d*_ = 0.1, Fig. 3), a strongly clustered spatial structure develops. This can be seen in individual realisations of the IBM (Fig. 3a) and values of pair correlation function greater than 1 for pairs less than distance *r* = 0.5 apart (Figure 3a). The pair correlation function drops below 1 for *r* > 0.5, indicating that the clusters are not randomly distributed, but are spaced regularly apart from one another. This is a consequence of competition making it difficult for any individual to survive in the neighbourhood surrounding an established cluster. These results are consistent with the translationally invariant version of the spatial stochastic logistic model. The cause of the clustering is the short dispersal distances leading to an accumulation of offspring around a common ancestor. This cluster eventually reaches a critical size where proliferation by individuals in the cluster is balanced by the elevated mortality rates due to competition. Because competition operates over a relatively long range, all individuals in a cluster tend to compete with all other individuals in the same cluster and hence experience similar mortality rates. Short-range dispersal makes it very difficult for new offspring to escape the cluster.

**Figure 3:**
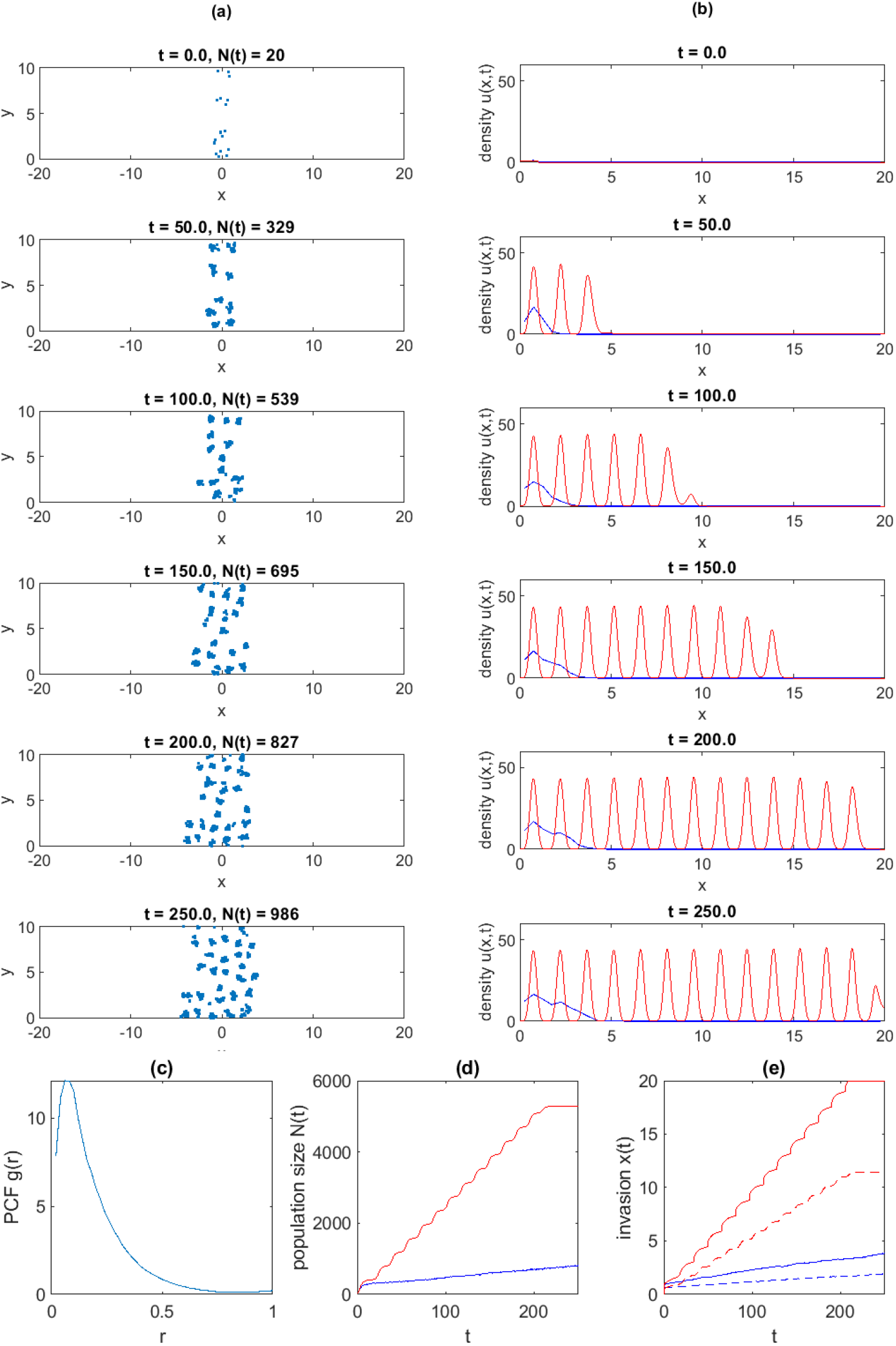
IBM and mean-field results for long-range competition and short-range dispersal (*σ*_*c*_ = 1, *σ*_*d*_ = 0.1): (a) snapshots of a single realisation of the IBM; (b) average agent density in the IBM (blue) and mean-field equation (red) at *t* = 0, 50, 100, 150, 200, 250. (c) pair-correlation function (PCF) at *t* = 250; (d) time series of the average population size in the IBM (blue) and mean-field equation (red); (e) time series of the invasion size measured by the location of the invasion front (solid) and the root mean squared displacement (dashed) in the IBM (blue) and mean-field equation (red). IBM results in (b-e) are averaged across *M* = 10 independent realisations, each initialised with *N*_0_ agents randomly placed in the region |*x*| < *x*_0_.

In the very early stages on the invasion up to around *t* = 20, the population size and invasion speed are reasonably well approximated by the mean-field equation (Fig. 3d,e). During this phase of the invasion, the population is increasing in density, but is mostly restricted to the initially occupied region, |*x*| ≤ *x*_0_. At around *t* = 20, the mean-field model establishes a newly occupied region and undergoes a second wave of rapid population growth. This pattern repeats periodically with the mean-field population alternating between phases of growth (increasing density in situ) and expansion (occupying new areas). In contrast, the clustered structure in the IBM makes it very difficult for a daughter agent to escape the influence of its ancestral cluster and establish a new cluster. Only occasionally can a new cluster establish and this means that the invasion proceeds very slowly relative to mean-field.

When both competition and dispersal and short-range (*σ*_*c*_ = *σ*_*d*_ = 0.1, Fig. 4), the spatial structure is also clustered, although not as strongly as when competition acts over a longer range (Fig. 3). Short-range dispersal means that individuals with a common ancestor have strongly correlated locations, but tend to be thinned out by short-range competition. Although the clustering is weaker than in Fig. 3, it still severely limits the ability of the population to invade, with population growth and the invasion speed much lower than predicted by the mean-field equation (Fig. 4d-e).

**Figure 4:**
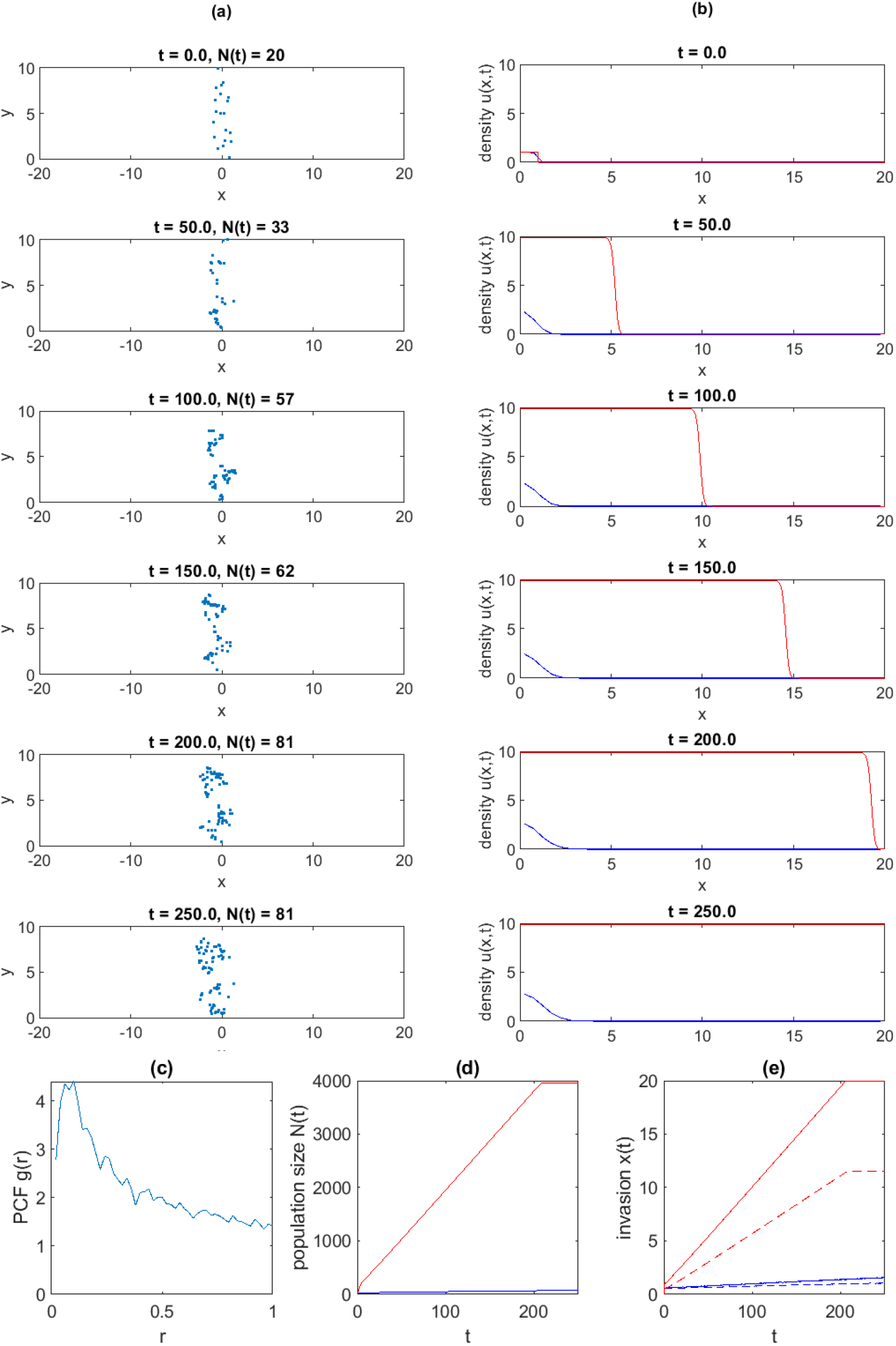
IBM and mean-field results for short-range competition and short-range dispersal (*σ*_*c*_ = 0.1, *σ*_*d*_ = 0.1): (a) snapshots of a single realisation of the IBM; (b) average agent density in the IBM (blue) and mean-field equation (red) at *t* = 0, 50, 100, 150, 200, 250.(c) pair-correlation function (PCF) at *t* = 250; (d) time series of the average population size in the IBM (blue) and mean-field equation (red); (e) time series of the invasion size measured by the location of the invasion front (solid) and the root mean squared displacement (dashed) in the IBM (blue) and mean-field equation (red). IBM results in (b-e) are averaged across *M* = 50 independent realisations, each initialised with *N*_0_ agents randomly placed in the region |*x*| < *x*_0_.

To understand the departures of the IBM from mean-field predictions, it is helpful to visualise how competition and dispersal interact spatially. Fig. 5 shows the strength of competition *C*(*x, y, t*) and the expected rate of arrival of newly dispersed offspring (i.e. propagule pressure) *P* (*x, y, t*), defined by

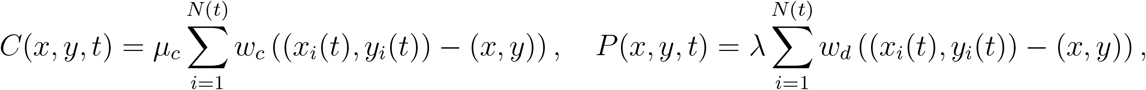

in a typical realisation of the IBM. When competition is short-range and dispersal is long-range (Fig. 5a-c), propagule pressure is uniformly high at all locations behind the invasion front (Fig. 5c). This means that any gaps in the patchy competition field (Fig. 5b) where an individual is viable will rapidly be filled. At the invasion front itself, there is a lower but still significant propagule pressure across the whole front, and so newly dispersed individuals can continuously advance the invasion front.

**Figure 5:**
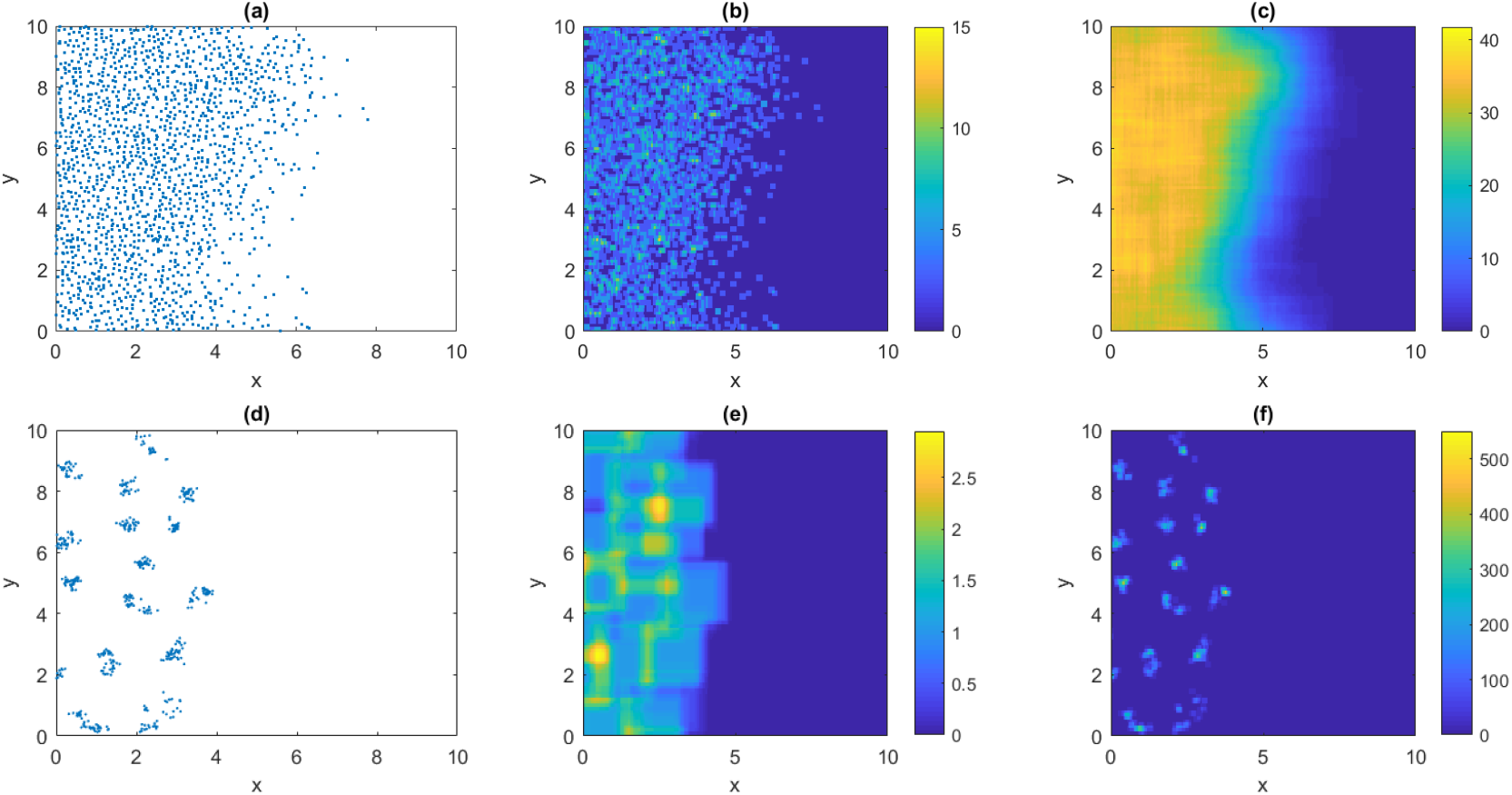
Maps of the strength of competition and dispersal in one typical realisation of the IBM with: (a-c) short-range competition and long-range dispersal (*σ*_*c*_ = 0.1, *σ*_*d*_ = 1); (d-f) long-range competition and short-range dispersal (*σ*_*c*_ = 1, *σ*_*d*_ = 0.1). (a,d) show agent locations; (b,e) show the strength of competition; (c,f) show the propagule pressure. Results in (a-c) correspond to the final time point of the IBM realisation shown in Fig. 2(a); results in (d-f) correspond to the final time point of the IBM realisation shown in Fig. 3(a).

When competition is long-range and dispersal is short-range (Fig. 5d-f), these spatial patterns are reversed: competition is strong everywhere behind the invasion front and at the front itself; propagule pressure is highly patchy and localised to the locations of existing clusters. This shows how difficult it is for the population to establish new clusters and hence advance the front.

We also tested the robustness of the model to the choice of interaction kernels by simulating the individual-based model and solving the mean-field equation using Gaussian kernels instead of the Heaviside kernels specified by Eq. (4). We also tested model sensitivity to the rate parameters λ, *μ*_0_ and *μ_c_*. Changing the kernels or parameter values did not qualitatively change the model behaviour or key results (see Supplementary Information).

## Discussion

The effect of spatial structure on average population density has been investigated previously (Law et al., 2003; Binny et al., 2016b). However, in some situations, understanding and predicting how spatial structure affects a biological invasion is more relevant than predictions of population density. Examples include the invasion of a pest species (Sprague et al., 2019), species range shifts due to climate change (Godsoe et al., 2014; Hurford et al., 2019), wound healing where cells migrate to fill injured tissue (Maini et al., 2004), or invasion of cancer cells into healthy tissue.

We have investigated the dynamics of translationally dependent populations under the spatial stochastic logistic model. This is a simple individual-based model (IBM) that consists of two mechanisms: density-independent proliferation accompanied by dispersal; and density-dependent mortality as a result of local competition (Bolker and Pacala, 1997; Law et al., 2003). Our results reveal that spatial structure can affect invasion speed and population density in different ways. In the long-range competition, short-range dispersal regime, the clustered spatial structure reduces both population density and invasion speed. The spatial structure in the occupied region rapidly reaches a strongly clustered state. This strong localised clustering makes it difficult for offspring to escape the competitive influence of their cluster and this is the limiting factor both for the effective carrying capacity and for the speed at which the invasion front can advance. In the short-range competition, long-range dispersal regime, a regular spatial structure develops. The uniformly high propagule pressure means that the population is effective at filling gaps in the competition field, and can therefore grow to higher densities than predicted by the mean-field. This is consistent with previous studies of the translationally invariant form of the model (Law et al., 2003). However, the speed of the invasion remains close to the mean-field prediction. This result may be understood by examining the spatial interaction of competition and dispersal (Fig. 5). Although short-range competition increases the effective carrying capacity in the interior of the population, the speed of the invasion is determined primarily by the low-density dynamics at the front itself. Thus, the limiting factors for the invasion speed are the net proliferation rate at low-density and the characteristic dispersal distance, as predicted by the mean-field model.

In all situations investigated, the IBM population invades at a similar or slower rate than the standard mean-field, suggesting that the mean-field equation provides an upper bound for the true invasion speed. The main factor that limits the invasion is the dispersal distance. Populations with short-range dispersal tend to invade more slowly, relative to mean-field predictions, because it is difficult for daughter agents to escape from the competitive influence of their ancestral cluster.

Lewis (2000) derived analytical and asymptotic results for the invasion speed in a one-dimensional, spatially structured population with competition and dispersal. These results, in contrast to ours, showed that short-range competition could significantly slow the asymptotic invasion speed. The discrepancy could be due to differences in model assumptions. In particular, the model of Lewis (2000) assumed that all individuals with at least one neighbour within a defined distance immediately became non-viable. In contrast, in our model, individuals in a crowded neighbourhood have an elevated probability of mortality. This means that crowded neighbourhoods tend to be thinned out over time, but some individuals will typically remain and continue to proliferate. Model behaviour in the low-density limit, which is the most determinant of invasion speed, is therefore less sensitive to competition than that of Lewis (2000). The results of Lewis (2000) were restricted to the short-range competition case so did not cover the clustered populations and reduced invasion speeds seen in our model under long-range competition.

The scenarios we have tested are similar to those investigated by Omelyan and Kozitsky (2019), who developed spatial moment dynamics approximation for a population with the same competition and dispersal mechanisms, described by Heaviside kernels, as used here. Omelyan and Kozitsky (2019) solved this system in one spatial dimension, but their solutions, particularly the speed of the invasion front, are yet to be tested against IBM simulations. Our individual-based simulations exhibit similar spatial structure to that predicted by Omelyan and Kozitsky (2019), i.e. clustered in the short-range dispersal regime and regular in the short-range competition regime. However, the spatial moment dynamics system of Omelyan and Kozitsky (2019) predicted that, in the short-range dispersal regime, the invasion speed would be similar to that of the mean-field equation and the population size would be larger. In contrast, our results for this case show that both the invasion speed and the population size in the IBM are much lower than mean-field. It is possible that differences between our results and those of Omelyan and Kozitsky (2019) are due to differences in the strength of spatial structure that develops in the one-dimensional and two-dimensional versions of the model, or the impact of approximations inherent in the analysis based upon a moment closure approximation (Murrell et al., 2004).

Our key results are robust to the exact shape of the dispersal and competition kernels, for example Heaviside or Gaussian kernels. However, we have not tested the effect of heavy-tailed kernels, which have been used to describe dispersal in empirical and theoretical studies (Kot et al., 1996; Katul et al., 2005; Paradis et al., 2002; Sundberg, 2005). It is likely that a heavy-tailed dispersal kernel would change the behaviour of the model because it means that, even when the majority of dispersal distance are small, there are occasional long-distance dispersal events. We hypothesise that this would improve the ability of offspring to escape the competitive influence of clusters and hence occupy new regions more rapidly, but this remains to be tested. It is known that heavy-tailed dispersal tends to lead to a patchy distribution without a well-defined invasion front (Lewis and Pacala, 2000) and an accelerating invasion that does not approach a constant speed (Kot et al., 1996). How these phenomena interact with small-scale spatial structure is an interesting question for future research.

We have focused on the spatial stochastic logistic model (Bolker and Pacala, 1997; Law et al., 2003), which is the simplest IBM that is capable of generating non-trivial spatial structure. There are other individual-level mechanisms that generate and/or are influenced by spatial structure. Examples include density-dependent proliferation (Lewis, 2000), movement (Dieckmann and Law, 2000; Murrell and Law, 2000), directional bias (Binny et al., 2015), and interspecific interactions (Bolker and Pacala, 1999; Murrell and Law, 2003). The interplay between these mechanisms and spatial structure has been investigated for translationally invariant populations, i.e. when the average density is spatially uniform (Binny et al., 2016b; Surendran et al., 2018; Binny et al., 2019). Extending the analysis of these mechanisms to a translationally dependent population will be an objective of future work.

## Acknowledgements

MP and RB acknowledge financial support from Te Pūnaha Matatini. MJS is supported by the Australian Research Council (DP170100474). We are grateful to Frederic Barraquand and one anonymous reviewer for critical comments on an earlier version of the manuscript.

