## Supplementary Information for "Small-scale spatial structure influences large-scale invasion rates"

### Supplementary results for Gaussian interaction kernels

To test the robustness of the model to different choices of interaction kernel, we performed individual-based model simulations and calculated mean-field solutions using Gaussian interaction kernels defined by

$$w_{[c,d]}(\xi) = \frac{1}{2\pi\sigma_{[c,d]}^2} \exp\left(-\frac{|\xi|^2}{2\sigma_{[c,d]}^2}\right) \quad \text{(S1)}$$

This replaces the Heaviside kernels defined by Eq. (4) in the main text.

Supplementary Figure S1 shows results in the case of short-range competition and long-range dispersal ($\sigma_c = 0.1$, $\sigma_d = 1$). Supplementary Figure S2 shows results in the case of long-range competition and short-range dispersal ($\sigma_c = 1$, $\sigma_d = 0.1$). All other parameter values as the same as in Table 1 of the main text.

The qualitative behaviour of the model is the same as when Heaviside kernels are used (compare Supplementary Figures S1 and S2 with Figures 2 and 3 respectively of the main text). In the case of short-range competition and long-range dispersal, the population develops a regular spatial structure, grows to a higher density in the interior of its range that predicted by the mean-field equation, but invades at a similar speed to the mean-field prediction. In the case of long-range competition and short-range dispersal, the population develops a clustered spatial structure and invades much more slowly than predicted by the mean-field equation.

### Parameter sensitivity

In the main text, we focused on the effect of the parameters $\sigma_c$ and $\sigma_d$, representing the spatial scale of competition and dispersal respectively. The model also has three rate parameters: the proliferation rate $\lambda$, the intrinsic death rate $\mu_0$ and the density-dependent death rate $\mu_c$. By a non-dimensionalisation of the time variable $t$ and the location variables $(x, y)$, these three parameters may be reduced to the dimensionless parameter ratio $\mu_0/\lambda$ .

The results in the main text are for $\mu_0/\lambda = 0.01$. This relatively small value of $\mu_0/\lambda$ means that density-dependence is the main source of mortality. Supplementary Figures S3 and S4 show results when $\mu_0/\lambda = 0.25$. The average density in the interior of the population is reduced by the increased intrinsic death rate but, again, the results are qualitatively similar to those shown in the main text.

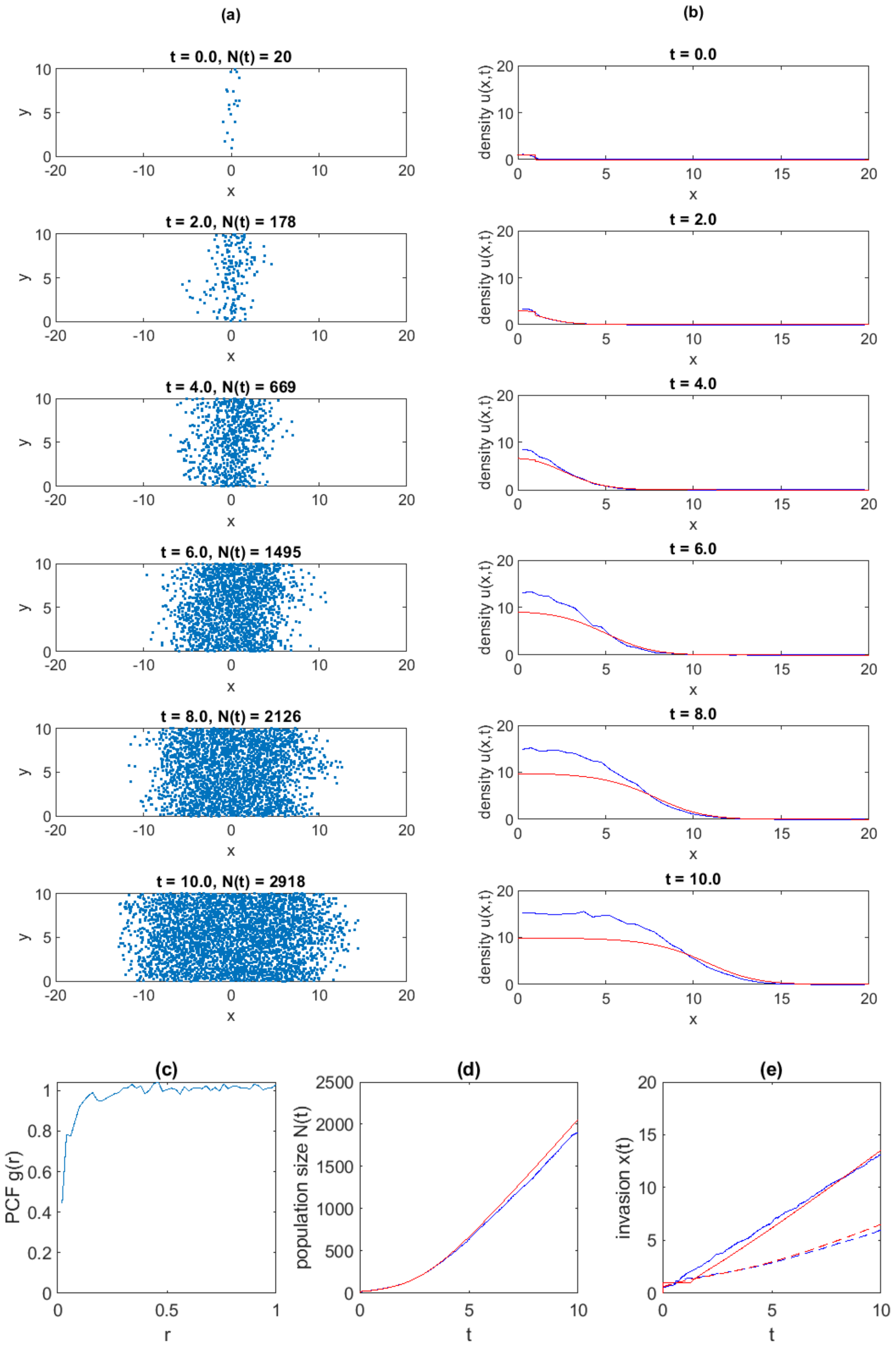


**Supplementary Figure S1**. IBM and mean-field results for Gaussian dispersal and competition kernels (Eq. S1) with short-range competition and long-range dispersal ($\sigma_c = 0.1$, $\sigma_d = 1$): (a) snapshots of a single realisation of the IBM; (b) average agent density in the IBM (blue) and mean-field equation (red) at $t = 0,2,4,6,8,10$. (c) pair correlation function (PCF) at $t = 10$; (d) time series of the average population size in the IBM (blue) and mean-field equation (red); (e) time series of the invasion size measured by the location of the invasion front (solid) and the root mean squared displacement (dashed) in the IBM (blue) and mean-field equation (red). IBM results in (b-e) are averaged across $M = 10$ independent realisations, each initialised with $N_0$ agents randomly placed in the region $|x| < x_0$. Other parameter values as shown in Table 1 of main text.

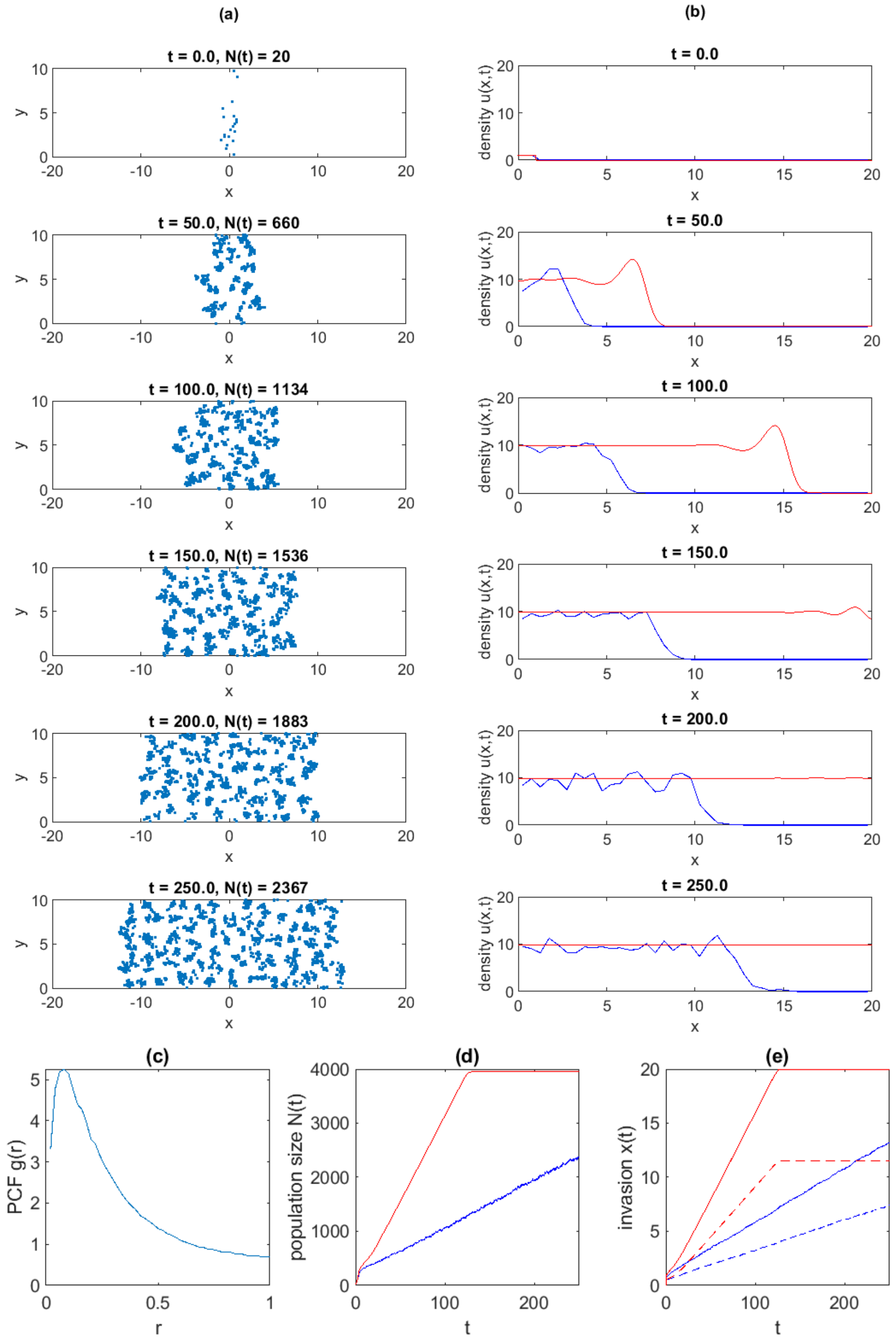


**Supplementary Figure S2**. IBM and mean-field results for Gaussian dispersal and competition kernels (Eq. S1) with long-range competition and short-range dispersal ($\sigma_c = 1, \sigma_d = 0.1$): (a) snapshots of a single realisation of the IBM; (b) average agent density in the IBM (blue) and mean-field equation (red) at $t = 0,50,100,150,200,250$. (c) pair correlation function (PCF) at $t = 250$; (d) time series of the average population size in the IBM (blue) and mean-field equation (red); (e) time series of the invasion size measured by the location of the invasion front (solid) and the root mean squared displacement (dashed) in the IBM (blue) and mean-field equation (red). IBM results in (b-e) are averaged across $M = 10$ independent realisations, each initialised with $N_0$ agents randomly placed in the region $|x| < x_0$. Other parameter values as shown in Table 1 of main text.

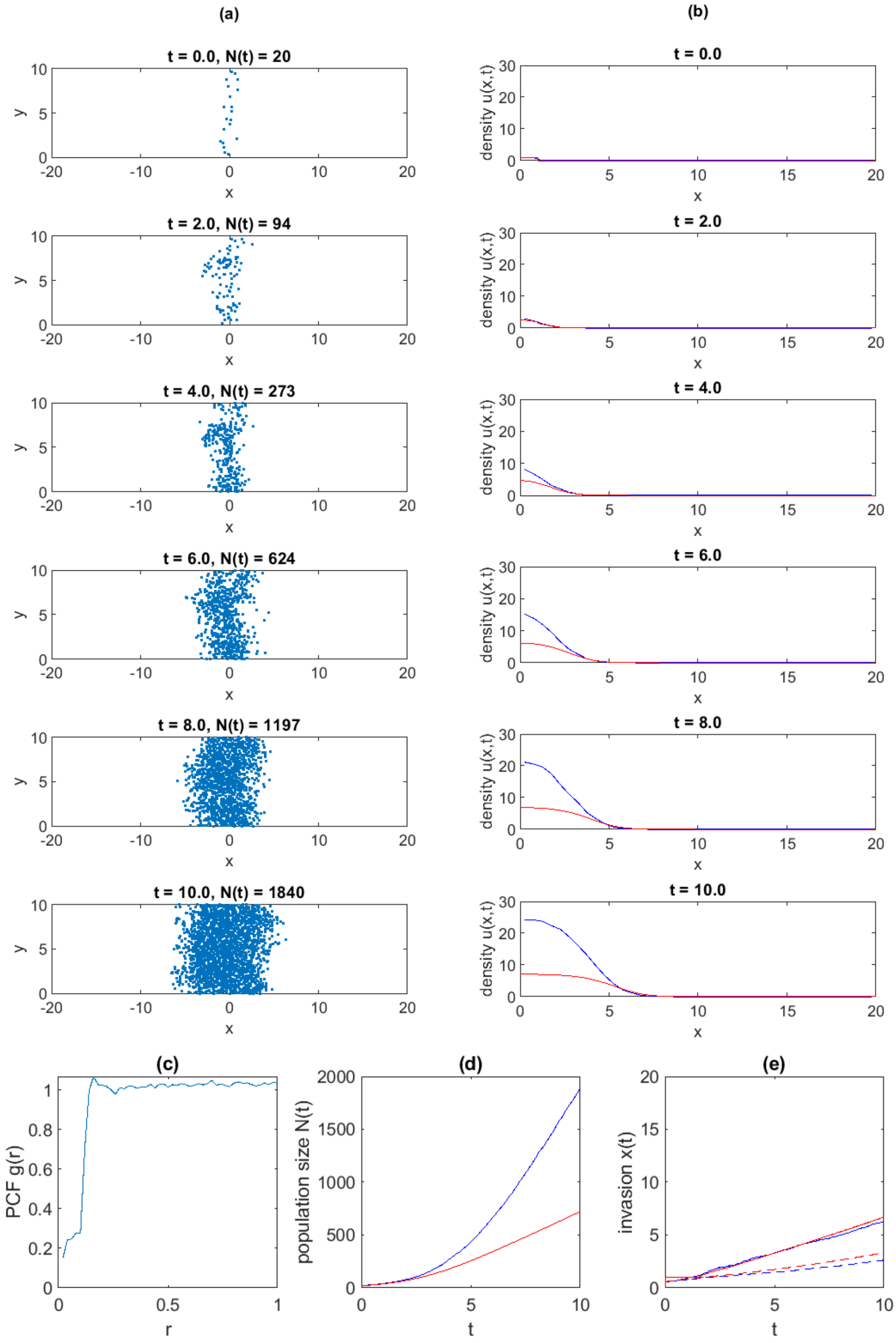


**Supplementary Figure S3**. IBM and mean-field results with increased intrinsic mortality rate ($\mu_0 = 0.25$), Heaviside dispersal and competition kernels, and short-range competition and long-range dispersal ($\sigma_c = 0.1$, $\sigma_d = 1$): (a) snapshots of a single realisation of the IBM; (b) average agent density in the IBM (blue) and mean-field equation (red) at $t = 0,2,4,6,8,10$. (c) pair correlation function (PCF) at $t = 10$; (d) time series of the average population size in the IBM (blue) and mean-field equation (red); (e) time series of the invasion size measured by the location of the invasion front (solid) and the root mean squared displacement (dashed) in the IBM (blue) and mean-field equation (red). IBM results in (b-e) are averaged across $M = 10$ independent realisations, each initialised with $N_0$ agents randomly placed in the region $|x| < x_0$. Other parameter values as shown in Table 1 of main text.

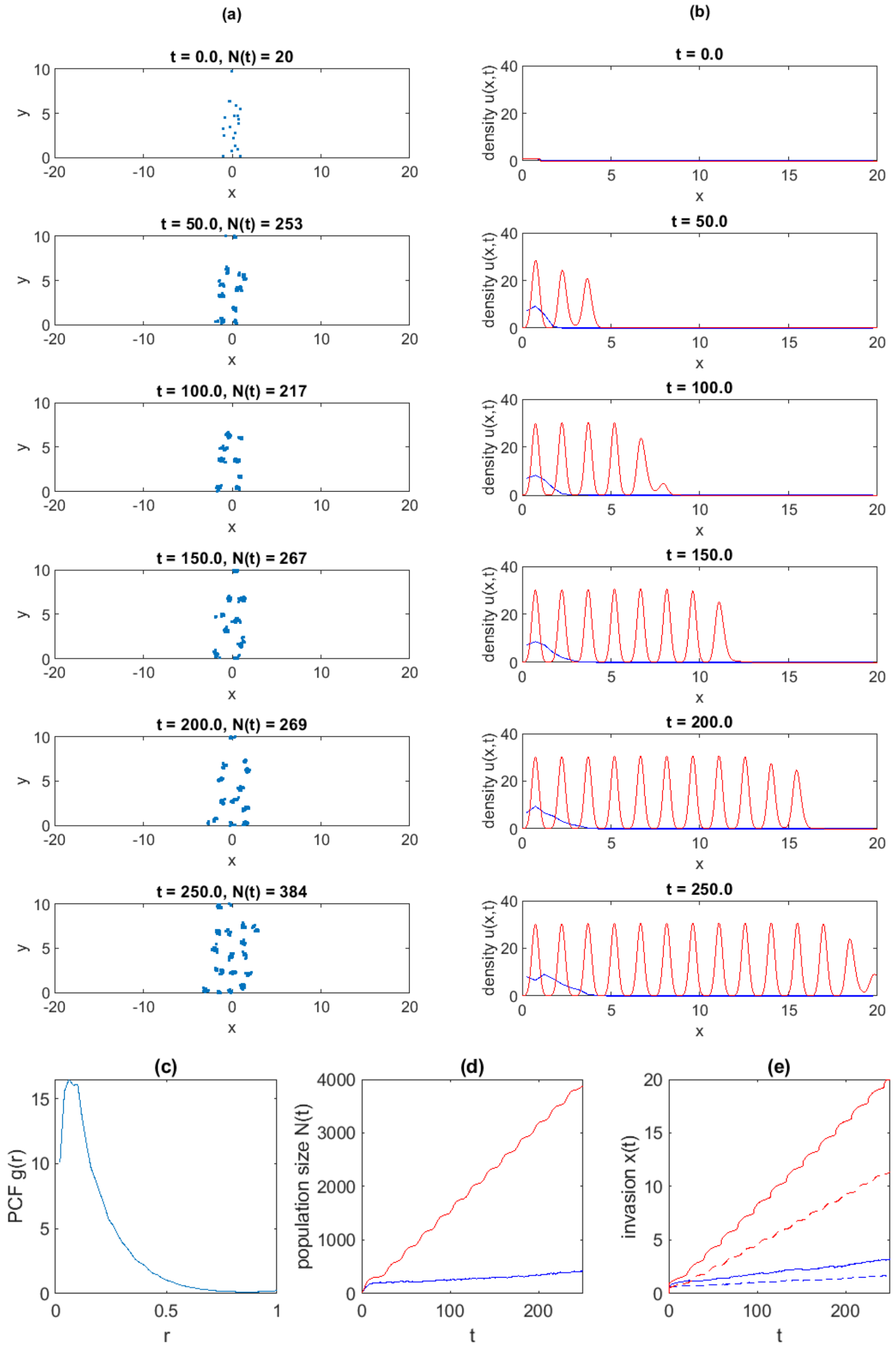


**Supplementary Figure S4**. IBM and mean-field results with increased intrinsic mortality rate ($\mu_0 = 0.25$), Heaviside dispersal and competition kernels, and long-range competition and short-range dispersal ($\sigma_c = 1$, $\sigma_d = 0.1$): (a) snapshots of a single realisation of the IBM; (b) average agent density in the IBM (blue) and mean-field equation (red) at $t = 0,50,100,150,200,250$. (c) pair correlation function (PCF) at $t = 250$; (d) time series of the average population size in the IBM (blue) and mean-field equation (red); (e) time series of the invasion size measured by the location of the invasion front (solid) and the root mean squared displacement (dashed) in the IBM (blue) and mean-field equation (red). IBM results in (b-e) are averaged across $M = 10$ independent realisations, each initialised with $N_0$ agents randomly placed in the region $|x| < x_0$. Other parameter values as shown in Table 1 of main text.
